# Microglial MHC-I induction with aging and Alzheimer’s is conserved in mouse models and humans

**DOI:** 10.1101/2023.03.07.531435

**Authors:** Collyn M. Kellogg, Kevin Pham, Adeline H. Machalinski, Hunter L. Porter, Harris E. Blankenship, Kyla Tooley, Michael B. Stout, Heather C. Rice, Amanda L. Sharpe, Michael J. Beckstead, Ana J. Chucair-Elliott, Sarah R. Ocañas, Willard M. Freeman

**Author notes:** These authors contributed equally (Collyn M. Kellogg, Kevin Pham). To whom correspondence should be addressed: Willard M. Freeman, Genes & Human Disease Program, Oklahoma Medical Research Foundation, 825 NE 13^th^ Street, Oklahoma City, OK 73104, USA.

## Abstract

Major Histocompatibility Complex I (MHC-I) CNS cellular localization and function is still being determined after previously being thought to be absent from the brain. MHC-I expression has been reported to increase with brain aging in mouse, rat, and human whole tissue analyses but the cellular localization was undetermined. Neuronal MHC-I is proposed to regulate developmental synapse elimination and tau pathology in Alzheimer’s disease (AD). Here we report that across newly generated and publicly available ribosomal profiling, cell sorting, and single-cell data, microglia are the primary source of classical and non-classical MHC-I in mice and humans. Translating Ribosome Affinity Purification-qPCR analysis of 3-6 and 18-22 month old (m.o.) mice revealed significant age-related microglial induction of MHC-I pathway genes *B2m*, *H2-D1*, *H2-K1*, *H2-M3*, *H2-Q6*, and *Tap1* but not in astrocytes and neurons. Across a timecourse (12-23 m.o.), microglial MHC-I gradually increased until 21 m.o. and then accelerated. MHC-I protein was enriched in microglia and increased with aging. Microglial expression, and absence in astrocytes and neurons, of MHC-I binding Leukocyte Immunoglobulin-like (Lilrs) and Paired immunoglobin-like type 2 (Pilrs) receptor families could enable cell-autonomous MHC-I signaling and increased with aging in mice and humans. Increased microglial MHC-I, Lilrs, and Pilrs were observed in multiple AD mouse models and human AD data across methods and studies. MHC-I expression correlated with *p16INK4A*, suggesting an association with cellular senescence. Conserved induction of MHC-I, Lilrs, and Pilrs with aging and AD opens the possibility of cell-autonomous MHC-I signaling to regulate microglial reactivation with aging and neurodegeneration.

## Introduction

Historically, the brain was proposed to be ‘immune-privileged’, due to the reported lack of/limited ability to develop adaptive immune responses to foreign antigens^1, 2^ and lack of conventional dendritic cells required to mount an adaptive immune response^3, 4^. In many cases, this difference between the brain and peripheral organ systems led to a presumption that immune-related proteins expressed in other organ systems had no or limited function in the normative CNS, except during viral infection and some pathological states where a highly compromised BBB allows leukocyte recruitment into the brain parenchyma to mount adaptive immune responses^5^. This has proven to be an overly simplistic assumption. Current molecular and biochemical studies have identified a number of immune-related pathways with normative CNS functions including, class I major histocompatibility complex (MHC-I)^6^, class II major histocompatibility complex (MHC-II)^7^, toll-like receptors (TLRs)^8^, and complement components^9, 10^. The characterization of these molecules as ‘immunological’ comes from their initial discovery in the immune system but these genes are functionally pleiotropic, and could be most broadly characterized as signaling molecules with different functions in different cellular milieus^11^. For example, mice deficient in complement components display defects in CNS synapse elimination^12, 13^, but this process is independent of the classical role of complement in innate immune clearance of microbes or adaptive immune detection of non-self antigens. Similarly, deficiency of the MHC-I molecule β2-microglobulin (*B2m*) leads to higher cortical synapse density in the mouse brain^14^. A more inclusive view of these ‘immunological’ molecules in the CNS is as signaling molecules that could work in both *cis* (cell-autonomous signaling within one cell) and *trans* (signaling between two cells, such as across synapses) to regulate brain function, in addition to conventional immunological roles. This fits with a more complete and nuanced contemporary view of immune privilege^15, 16^.

The MHC pathways are elements of the adaptive and innate immune systems, with Class I canonically presenting intracellular antigens for viral surveillance and Class II presenting extracellular antigens for bacterial surveillance. While canonically thought to be unexpressed in the brain^17, 18^, evidence of MHC-I expression by brain resident cells, principally glia, has been shown for decades^19, 20^. MHC-I is further divided into classical (I/Ia) and non-classical (Ib) groupings in which the non-classical genes exhibit a more restricted expression to specific cell types and are less polymorphic^21^. There are no exact homologs between each murine and human MHC-I genes but Class I in humans encompasses *HLA-A*, *-B*, and *-C*, and in Mice *H2-D* and *-K* forms, while Class Ib in humans are *HLA-E*, *-F* and *-G,* and in mice are *H2-M*, *-T*, and *-Q*^22, 23^. Peptides for presentation by MHCs are generated initially in the cytosol by the proteasome^24^. These peptides are transported into the endoplasmic reticulum (ER) by the transporter associated with antigen processing (*Tap1* and *Tap2*). Peptides are then loaded onto MHCs by tapasin (*Tapbp*, transporter associated with antigen processing binding protein). MHC-I can also cross-present exogenous antigens^25^. For MHCs to localize to the cell surface, they must be associated with *B2m*^26^. MHC-I is canonically recognized by the T-Cell receptors (TCRs) to mount adaptive immune responses in an antigen-dependent manner^27^. However, a wide variety of paired immune receptors can also bind MHC-I in what is likely antigen-dependent and antigen-independent manners (though many questions remain to be answered in this signaling process)^28–30^. These paired receptor families include leukocyte immunoglobulin-like receptors (Lilrs)^31, 32^, paired immunoglobin-like type 2 receptors (Pilrs)^33^, and C-type lectin-like receptors (Clecs)^34^, most of which contain ITAMs (Immunoreceptor Tyrosine-based Activation Motifs) and ITIMs (Immunoreceptor Tyrosine-based Inhibition Motifs) that can induce activational or inhibitory signals in the recognizing cell^35^.

After first being described as absent from the CNS^17^, especially in neurons^18^, CNS MHC-I expression was ascribed to neurons in developmental studies^20, 36^ and then in the adult but without expression in microglia or astrocytes^37, 38^, except in pathological states. Much of the interest in MHC-I and potential non-TCR mediated signaling came from developmental literature^39–42^ and proposed a neuronal source and function of MHC-I. Recently, a neuron-centric MHC-I function in AD was reported^43^ where MHC-I, driven by ApoE expression, induces neurodegeneration. Other studies have ascribed MHC-I expression to astrocytes^44^, microglia^45^, and other CNS cell types^46^. However, microglial molecular phenotypes developed from scRNA-Seq studies include MHC-I components and paired receptor family members^47–49^ as key markers of phenotypic states including Disease Associated Microglia (DAMs) and other subtypes (recognizing that the microglial state terminology has become overlapping and occasionally misleading^50^). MHC-I components have also been proposed as markers of astrocyte phenotypes^51^. This lack of clarity of which CNS cells expression MHC-I components may arise from these studies examining a variety of ages, pathological states, and different brain regions and the different methods employed that each have their own strengths and limitations.

Previously, we have reported MHC-I induction with aging in hippocampal dissections of mice and rats^52, 53^, and this study seeks to clarify the cell type(s) expressing MHC-I in the brain and localize the cellular source of MHC-I with aging and AD, in mice and humans. Data across multiple cohorts of animals/subjects and from a variety of methods were combined to increase the rigor of analyses and avoid technical confounds that may be the source of the disparate findings in the literature. Microglia were identified as the primary source of MHC-I components (though this does not imply absolute restriction to microglia) and were the source of increased expression with aging and AD. Co-expression of MHC-I with paired immunoglobulin-like receptors (Lilr and Pilr gene families) that can bind MHC-I opens the possibility of cell-autonomous signaling of MHC-I to these receptors and regulation of microglial phenotype by MHC-I with aging and neurodegenerative disease. Furthermore, microglial MHC-I was found to follow the same timecourse and correlate to markers of cellular senescence with aging and AD.

## Methods

*Mice:* All animal procedures were approved by the Oklahoma Medical Research Foundation (OMRF). Mice were purchased from the Jackson Laboratory (Bar Harbor, ME), bred, and housed at the OMRF, under SPF conditions in a HEPA barrier environment. In separate breeding strategies, as described previously^54^, Aldh1l1-Cre/ERT2^+/wt^ (stock number # 031008)^55^, Cx3cr1^Jung^-Cre/ERT2^+/+^ (stock # 20940)^56^, Cx3cr1^Litt^-Cre/ERT2^+/+^ (stock #021160)^57^, and CamkIIα-Cre/ERT2^+/wt^ (stock # 012362)^58^ males were mated with NuTRAP^flox/flox^ females (stock # 029899)^59^ to generate the desired progeny, Aldh1l1-cre/ERT2^+/wt^; NuTRAP^flox/wt^ (Aldh1l1-NuTRAP), Cx3cr1^Jung^-cre/ERT2^+/wt^; NuTRAP^flox/wt^ (Cx3cr1^Jung^-NuTRAP), Cx3cr1^Litt^-cre/ERT2^+/wt^; NuTRAP^flox/wt^ (Cx3cr1^Litt^-NuTRAP), and CamkIIα-cre/ERT2^+/wt^; NuTRAP^flox/wt^ (CamkIIα-NuTRAP). Founders for both APP-PSEN1 (MMRC #034832)^60^ and control mice were obtained from Dr. Salvatore Oddo, who had backcrossed them for more than 12 generations and maintained them on a 129/SvJ background as previously described^61^. Local breeding of APP-PSEN1 and control mice was done in parallel. At 3 m.o. in the NuTRAP models, unless otherwise noted, mice received a daily intraperitoneal (ip) injection of tamoxifen (Tam) solubilized in 100% sunflower seed oil by sonication (100□mg/kg body weight, 20□mg/ml stock solution, #T5648; Millipore Sigma, St. Louis, MO) for five consecutive days^54^. TAM dependent Cre induction at 3 m.o. does not lead to long-lasting changes in brain gene expression^62^ and avoids potential confounds of early post-natal TAM administration^63^. Based on an average weight of 20 g per mouse, each daily injection of Tam consisted of 100□μl of 20□mg/ml stock solution. Adjustments were made for mice that significantly deviated from the average weight. For TAM induced NuTRAP mice at least three weeks passed since last TAM injection before sample collection. In Cx3cr1-NuTRAP mice this allows clearance of any labeled circulating Cx3cr1^+^ cells. All aged mice were induced at 3 m.o. as well. Mice were euthanized by cervical dislocation, followed by rapid decapitation, in line with the AVMA Guidelines for the Euthanasia of Animals.

*Mouse Genotyping*: DNA was extracted from mouse ear punch samples for genotyping by incubating samples in 50 nM NaOH at 95°C for 1 hour, after which the reaction was neutralized by adding 30 uL 1 M Tris HCl (pH: 7.4). Processed samples were then genotyped using a touchdown PCR reaction (94°C hotstart for 2 min, 10 cycles of touchdown (94°C 20 sec, 65°C 15 sec (–0.5C per cycle decrease per cycle)), 68°C,10 sec) followed by 28 cycles of amplification (94°C 15 sec, 60°C 15 sec, 72°C 10 sec) with the listed primer sets (**Supplemental Table 1**).

*TRAP isolation:* TRAP isolation from whole brain or hippocampus, as noted, was performed as previously described^54^. Minced tissue was Dounce homogenized (#D8938; MilliporeSigma) in 1.5 ml chilled homogenization buffer (50 mm Tris, pH 7.4; 12 mm MgCl_2_; 100 mm KCl; 1% NP-40; 1□mg/ml sodium heparin; 1 mm DTT; 100□μg/ml cycloheximide (#C4859-1ML, MilliporeSigma); 200 units/ml RNaseOUT Recombinant Ribonuclease Inhibitor (#10777019; Thermo Fisher Scientific); 0.5 mm Spermidine (#S2626, MilliporeSigma); 1× complete EDTA-free Protease Inhibitor Cocktail (#11836170001; MilliporeSigma)). Homogenates were transferred to 2 ml round-bottom tubes and centrifuged at 12,000 × *g* for 10□min at 4°C. 100□μl of supernatant was saved as “Input.” The remaining supernatant was transferred to a 2-ml round-bottom tube and incubated with 5 μg/μl of anti-GFP antibody (ab290; Abcam) at 4°C with end-over-end rotation for 1 h. Dynabeads Protein G for IP (#10003D; Thermo Fisher Scientific) were pre-washed three times in 1-ml ice-cold low-salt wash buffer (50 mm Tris, pH 7.5; 12 mm MgCl_2_; 100 mm KCl; 1% NP-40; 100□μg/ml cycloheximide; 1 mm DTT). The homogenate/antibody mixture was transferred to the 2-ml round-bottom tube containing the washed Protein-G Dynabeads and incubated at 4°C with end-over-end rotation overnight. Magnetic beads were collected (DynaMag-2 magnet) and the unbound-ribosomes and associated RNA discarded. Beads and GFP-bound polysomes were then washed 3X with 0.5 ml of high-salt wash buffer (50 mm Tris, pH 7.5; 12 mm MgCl_2_; 300 mm KCl; 1% NP-40; 100□μg/ml cycloheximide; 2 mm DTT). Following the last wash, 350□μl of buffer RLT (QIAGEN) supplemented with 3.5 μl 2-β mercaptoethanol (#444203, MilliporeSigma) was added directly to the beads and incubated with mixing on a ThermoMixer (Eppendorf) for 10□min at room temperature. The beads were magnetically separated and the supernatant containing the target bead-bound polysomes and associated RNA was transferred to a new tube and constitutes the “TRAP” or positive fraction for subsequent analysis. A total of 350□μl of 100% ethanol was added to the sample and loaded onto a RNeasy MinElute column (QIAGEN). RNA was isolated using RNeasy Mini kit (#74104, QIAGEN), and quantified with a Nanodrop One^c^ spectrophotometer (#ND-ONEC-W, Thermo Fisher Scientific) and quality assessed by HSRNA ScreenTape (#5067-5579, Agilent Technologies) with a 4150 Tapestation analyzer (#G2992AA, Agilent Technologies).

*RT-qPCR:* Relative gene expression levels were quantified by qPCR as previously described^64, 65^. cDNA was synthesized with the ABI High-Capacity cDNA Reverse Transcription Kit (Applied Biosystems Inc., Foster City, CA) from 25 ng of purified RNA. qPCR was performed with gene-specific primer probe fluorogenic exonuclease assays (TaqMan, Life Technologies, Waltham, MA, **Supplemental Table 2**) on a QuantStudio 5 Real-Time PCR System (Applied Biosystems). Relative gene expression (RQ) was calculated with Expression Suite v 1.3 software using the 2^−ΔΔ^Ct analysis method with GAPDH as an endogenous control. Statistical analysis of the qPCR data was performed using GraphPad Prism 9 (San Diego, CA). One or Two-way ANOVA analyses were performed as appropriate followed by the Tukey’s multiple comparison test (*p*L<L0.05) with a Benjamini-Hochberg Multiple Testing correction applied within each cell type or tissue.

*Single cell suspension:* Hippocampi were rinsed in ice-cold D-PBS containing calcium, magnesium, glucose, and pyruvate (#14287-072, Thermo Fisher Scientific), sliced into four sagittal sections on a chilled, metal block and placed into ice-cold gentleMACS C-tubes (#130-093-237, Miltenyi Biotec), containing 1950□μl of papain-based Enzyme Mix 1. For each reaction, Enzyme Mix 1 was created by combining 50□μl of Enzyme P with 1900□μl of buffer Z, while Enzyme Mix 2 was created by combining 10□μl of Enzyme A with 20□μl of buffer Y per reaction, with all reagents included in the Adult Brain Dissociation kit (#130-107-677, Miltenyi Biotec). Transcription and translation inhibitors were included during cell preparation to prevent *ex vivo* activational artifacts, as previously described^66^. Actinomycin D (#A1410, MilliporeSigma) was reconstituted in DMSO at 5Lmg/ml before being aliquoted and stored at −20°C protected from light. Triptolide (#T3652, MilliporeSigma) was reconstituted in DMSO to a concentration of 10 mM before being aliquoted and stored at −20°C protected from light. Anisomycin (#A9789, MilliporeSigma) was reconstituted in DMSO to a concentration of 10□mg/ml before being aliquoted and stored at 4°C protected from light. 2 μl each of actinomycin D, triptolide, and anisomycin stocks were added to the initial Enzyme Mix 1 before dissociation for a final concentration of 5 μg/ml, 10 μm, and 10□μg/ml, respectively. Each sample had 30□μl of Enzyme Mix 2 added before being mechanically dissociated for 30□min at 37°C on the gentleMACS Octo Dissociator with Heaters (#130-096-427, Miltenyi Biotec) using the 37C_ABDK_02 program. Following enzymatic and mechanical dissociation, the C-tubes were quickly spun in a chilled (4°C) Allegra-30R centrifuge (#B08708, Beckman Coulter) with an SX4400 swinging bucket rotor to collect the samples in the bottom of the tubes. Next, samples were resuspended, passed through a pre-wet 70□μm MACS SmartStrainer (#130-110-916, Miltenyi Biotec) and collected in a 50-ml conical tube (#21008-178, VWR International). The C-tubes were washed with 10 ml of ice-cold D-PBS and the washed volume was passed through the 70□μm MACS SmartStrainer. The cells were then pelleted by centrifugation at 300 × *g* for 10□min at 4°C. Following centrifugation, the supernatant was aspirated and debris was removed using the Debris Removal solution (#130-109-398, Miltenyi Biotec) provided in the Adult Brain Dissociation kit (#130-107-677, Miltenyi Biotec). Briefly, cells were resuspended in 1.55 ml of ice-cold D-PBS, and passed to a 5-ml round bottom tube (#22171606, FisherScientific) and 450□μl of cold Debris Removal solution was mixed into the cell suspensions. Next, 2 ml of D-PBS was gently overlaid on the cell suspension, ensuring the layers did not mix. Centrifugation at 3000 × *g* for 10□min at 4°C separated the suspension into three phases, of which the top two phases were aspirated. The cell pellet was gently resuspended in 5 ml of ice-cold D-PBS before centrifugation at 1000 × *g* for 10□min at 4°C. After aspirating the supernatant completely, the cells were resuspended in 1 ml 0.5% BSA (#130-091-376, Miltenyi Biotec) in D-PBS and filtered through a 35-μm filter (#352235, Fisher Scientific). A 100 μl aliquot of cells was retained as “Cell-Input” for comparison^66^.

*Microglial isolation – AutoMACS:* Hippocampal single cell suspensions were pelleted at 300 × g for 10□min at 4°C and resuspended in 90□μl of 0.5% BSA in D-PBS with 10□μl of CD11b (Microglia) MicroBeads (#130-093-636, Miltenyi Biotec) per 10^7^ total cells. After gentle pipette mixing, cells were incubated for 15 min at 4°C in the dark. Cells were washed with 1 ml of 0.5% BSA in D-PBS and pelleted at 300 × g for 10□min at 4°C. The cell pellet was resuspended in 500□μl of 0.5% BSA in D-PBS. After priming the autoMACS Pro Separator (#130-092-545, Miltenyi Biotec), sample and collection tubes were placed in a cold MACS Chill 5 Rack (#130-092-951, Miltenyi Biotec) with both positive and negative fractions being collected. The double-positive selection (Posseld) program (i.e., positive fraction cells are then run over a second magnetic column for higher purity) was used to elute highly pure CD11b+ cells in 500□μl of autoMACS Running buffer (#130-091-221, Miltenyi Biotec). Following separation, the positive and negative fractions were reserved for further analysis.

*RNA-Seq:* Directional RNA-Seq libraries (NEBNext Ultra II Directional RNA Library, New England Biolabs, Ipswich, MA NEB#E7760) were made according to the manufacturer’s protocol for 2 to 100□ng RNA as previously^66^. CD11b^+^ cells from magnetic bead isolation were used to create individual RNA-Seq libraries (no pooling of samples was performed). Poly-adenylated RNA was captured using NEBNext Poly(A) mRNA Magnetic Isolation Module (#NEBE7490). Following mRNA capture, mRNA was eluted from the oligo-dT beads and fragmented by incubating with the First Strand Synthesis Reaction buffer and Random Primer Mix (2×) from the NEBNext Ultra II Directional Library Prep Kit for Illumina (#NEBE7760; New England Biolabs) for 15 min at 94°C. First and second strand cDNA synthesis were performed sequentially, as instructed by the manufacturer’s guidelines. After purification of double-stranded cDNA with 1.8× SPRISelect Beads (#B23318, Beckman Coulter), purified cDNA was eluted in 50□μl 0.1× TE buffer and subjected to end prep. The NEBNext adaptor was diluted 1:100 in Adaptor Dilution buffer (provided) before ligating the adaptor to the cDNA. After purifying the ligation reaction with 0.9× SPRISelect Beads (#B23318, Beckman Coulter), cDNA was eluted in 15 μl of 0.1× TE (provided). Next, cDNA libraries were amplified with 16 cycles of PCR using the NEBNext Ultra II Q5 Master Mix (provided) and unique index primer mixes from NEBNext Multiplex Oligos for Illumina Library (#E6609L, New England Biolabs). Libraries were purified with 0.9× SPRISelect Beads (#B23318, Beckman Coulter) and then sized with HSD1000 ScreenTapes (#5067-5584; Agilent Technologies). Libraries had an average peak size of 316 bp. Libraries were quantified by Qubit 1× dsDNA HS Assay kit (#Q33230, Thermo Fisher Scientific). The libraries for each sample were pooled at 4 nm concentration and sequenced using an Illumina NovaSeq 6000 system (S4 PE150). The entirety of the sequencing data are available for download in FASTQ format from NCBI Gene Expression Omnibus (to be included on acceptance).

*Flow cytometry:* For flow cytometric analysis, cell preparations were taken for analysis on the MACSQuant Analyzer 10 Flow Cytometer. Cell were stained with CD11b-APC (M1/70, #130-113-793, Miltenyi Biotec) and CD45-VioBlue (REA737, #130-110-802, Miltenyi Biotec) and either H-2-PE (REA857, #130-112-481) or H-2-FITC (M1/42, #125508). Following staining, cells were resuspended in 250□μl of 0.5% BSA/D-PBS and run on the MACSQuant Analyzer 10 Flow Cytometer. Data were analyzed using MACSQuantify v2.13.0 software.

*Immunohistochemistry:* Brain samples were fixed for 4 h in 4% PFA, cryoprotected by sequential incubations in PBS containing 15% and 30% sucrose, and then frozen in Optimal Cutting Temperature medium (#4583, Tissue-Tek). Twelve μm-thick sagittal sections were cryotome-cut (Cryostar NX70, Thermo Fisher Scientific). Tissue sections were rinsed with PBS containing 1% Triton X-100, blocked for 1 h in PBS containing 10% normal donkey serum, and processed for fluorescence immunostaining and downstream analysis. The primary antibodies included rat anti-CD11b (#C227, Clone M1/70, 1:100, Leinco Technologies),rabbit anti-B2m (#ab75853, lot: GR237909-26, 1:100, Abcam), rabbit anti-NeuN (#ab177487, 1:200, lot: GR3250076-3, Abcam), mouse anti-MHC Class I (H-2Db) (#14-5999-85, 1:100, clone: 28-14-8, eBioscience), and mouse anti-MHC Class I RT1A (#MCA51G, 1:100, clone: OX-18, 1:100, Bio-Rad). Sequential imaging was performed on an Olympus FluoView confocal laser-scanning microscope (FV1200; Olympus) at the Dean McGee Eye Institute imaging core facility at OUHSC. Microscope and FLUOVIEW FV1000 version 1.2.6.0 software (Olympus) settings were identical and at the same magnification within a given experiment. The experimental format files were.oib. The Z-stack generated was achieved at 1.00-μm step size with a total of three optical slices (Figure 5E) and a total of nine optical slices (Figure 5A-D) at 40× magnification (2-2.5× zoom).

*Data Analysis:* Following sequencing, reads were trimmed and aligned before differential expression statistics and correlation analyses in Strand NGS software package (v4.0; Strand Life Sciences). Reads were aligned against the full mm10 genome build (2014.11.26). Alignment and filtering criteria included the following: adapter trimming, fixed 2-bp trim from 5′ and 2Lbp from 3′ ends, a maximum number of one novel splice allowed per read, a minimum of 90% identity with the reference sequence, a maximum 5% gap, and trimming of 3′ ends with QL<L30. Alignment was performed directionally with Read 1 aligned in reverse and Read 2 in forward orientation. All duplicate reads were then removed. Normalization was performed with the DESeq2 algorithm. Transcripts with an average read count value >5 in at least 100% of the samples in at least one group were considered expressed at a level sufficient for quantitation per tissue and those transcripts below this level were considered not detected/not expressed and excluded, as these low levels of reads are close to background and are highly variable. For statistical analysis of differential expression, a t-test between control and APP-PSEN1 with Benjamini–Hochberg multiple testing correction (BHMTC) was performed. For those transcripts meeting this statistical criterion, a fold change >|1.25| cutoff was used to eliminate those genes which were statistically significant but unlikely to be biologically significant and orthogonally confirmable because of their very small magnitude of change. Visualizations of hierarchical clustering and principal component analyses (PCAs) were performed in Strand NGS (version 4.0). Upset plots were created using ComplexHeatmap v2.14.0 package in RStudio 2022.07.2 Build 576 with R v4.2.1. Gene expression data and associated annotations of age and tissue were downloaded in bulk from the Tabula Muris^67^ or Atlas of the Aging Mouse Brain^68^. Only counts greater than 1 were considered for the plot. For human GeTX data across tissues and ages, count-level expression data were quantile normalized and log-transformed before analysis as previously^52^. Significance of age-association was determined using Pearson correlation computed using Scipy stats’ pearsonr function^69^.

## Results

*MHC-I cellular expression in mice:* To determine the expression of MHC-I in murine CNS cell types RNA-Seq data from two different isolation methodologies, cell sorting and TRAP isolations was collected from public repositories. RNA-Seq data from bulk FACS or immunopanned young (∼2 m.o.) sorted astrocytes (Aldh1l1+), neurons (L1cam+), and microglia (CD45+) from the BrainRNASeq^70^ database were examined for Class 1, Class 1b, and antigen processing components of the MHC-I pathway. Limited to no expression was observed in astrocytes or neurons, with highest levels of expression in microglia (**Figure 1A, Supplemental Table 3**). Cell type-specific translatome RNA-Seq data from Translating Ribosome Affinity Purification (TRAP)^71^ isolated hippocampal RNA from young (∼3 m.o.) mice were compared. Three previously validated mouse lines^54^ for astrocytes (Aldh1l1-NuTRAP), neurons (CamkIIα-NuTRAP), and microglia (Cx3cr1-NuTRAP) were used (**Figure 1B, Supplemental Table 4**). Again, highest expression was observed in microglia. A difference between sorted cells and ribosomal profiling data is that transcripts translated in distal cellular processes are retained in TRAP isolates while these are generally lost in cell sorting data. Data comparing soma enriched and distal process enriched microglial transcripts^72^ identified B2m, H2-D1, H2-K1, and H2-M3 as all significantly enriched in distal processes of microglia.

**Figure 1:**
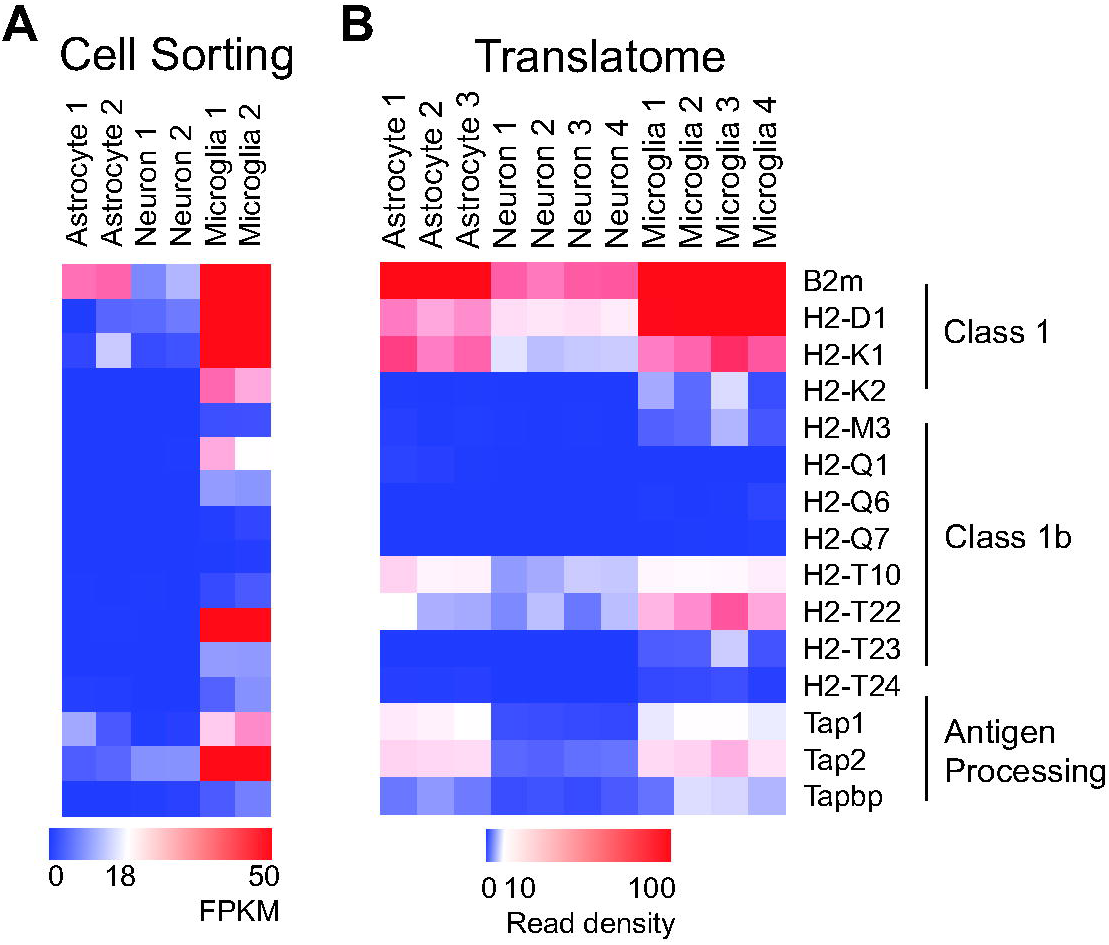
MHC-I pathway expression in mouse CNS cell types. Using bulk RNA-Seq data from **A)** sorted CNS cell types (∼2 m.o.) and from **B)** TRAP-Seq data (∼3 m.o.) of cell type-specific NuTRAP mice lines, α and β chains of MHC-I and Tap complex components are most highly expressed in microglia with minimal to no expression in astrocytes and neurons.

*Cell type-specific analysis of mouse hippocampal aging:* To examine hippocampal expression of MHC-I pathway components with aging in a cell type-specific manner, TRAP isolated RNA from our NuTRAP models for astrocytes (Aldh1l1-NuTRAP), neurons (CamkIIα-NuTRAP), and microglia (Cx3cr1-NuTRAP) at young (3-6 m.o.) and old (18-22 m.o.) ages, and from male and female mice was analyzed by RT-qPCR. Genes were selected based on existing tissue level RNA-Seq and scRNA-Seq analyses of mouse brain aging. Robust increases in *B2m*, as well as Class I (*H2-D1*, *H2-K1*) and non-classical Class Ib (*H2-M3*, *H2-Q6*) MHCs, and Tap complex member *Tap1* expression were observed in microglia, in both males and females with age (**Figure 2A, Supplemental Table 5**). A few, small magnitude age and sex differences were observed in astrocytes and neurons but no consistent regulation with aging was observed (**Figure 2B & C**).

**Figure 2:**
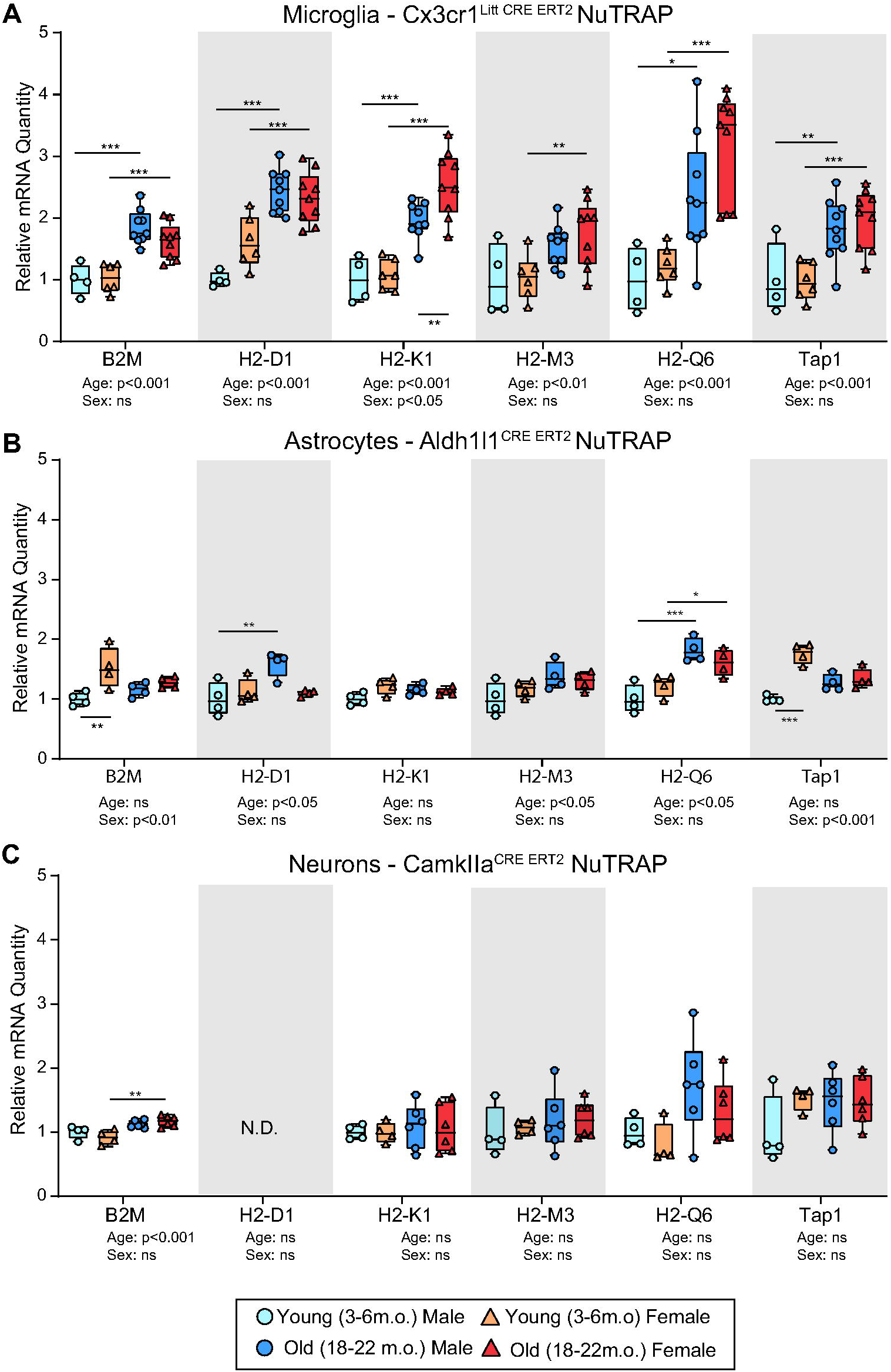
Microglial MHC-I expression induction with aging but not in other cell types. **A)** Cell type-specific hippocampal RNA from microglial (Cx3cr1^Litt^-NuTRAP), **B)** astrocytes (Aldh1l1-NuTRAP), and **C)** neurons (CamkIIα-NuTRAP) was isolated by TRAP. qPCR analysis was performed on Young (3-6m.o) and Old (18-22m.o) male and female mice. Increased microglial levels of MHC-I α and β chains and Tap1 were evident in males and females with aging. Minimal or no changes with age were evident in astrocytes and neurons. Two-Way ANOVA (Factors: Age and Sex) with SNK post hoc comparisons (*p<0.05, **p<0.01, ***p<0.001).

*Trajectory of mouse hippocampal MHC-I analysis with aging:* To more finely determined the age at which microglial MHC-I induction occurs, a timecourse of MHC-I pathway induction was examined across 12 – 23 m.o. in female Cx3cr1^Litt^-NuTRAP mice by TRAP-qPCR. A steady increase in MHC-I pathway expression was observed across all genes, with the largest increases most notably starting at 21 months of age (**Figure 3, Supplemental Table 6**). These data demonstrate a reproducible increase in microglial MHC-I with aging that accelerates as mice near two years of age.

**Figure 3:**
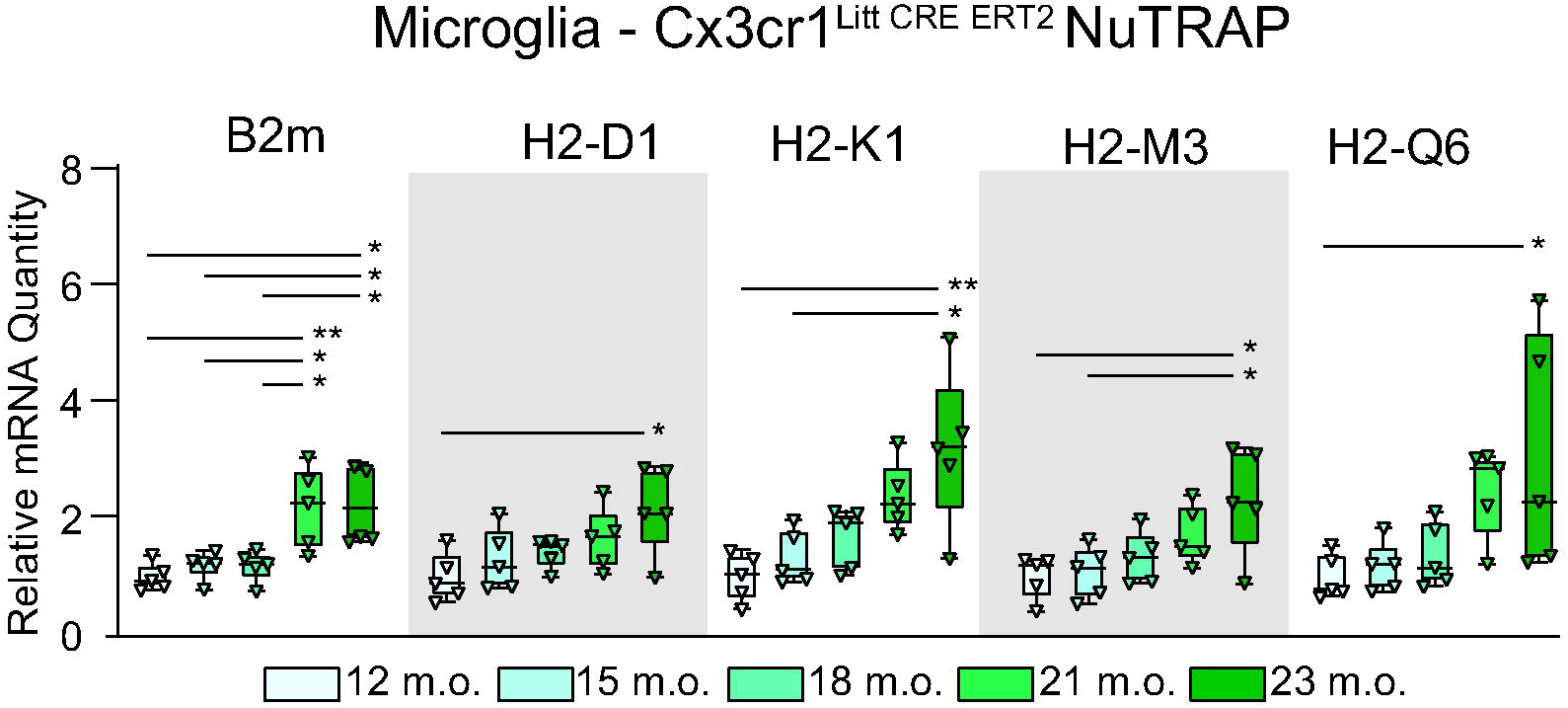
Timecourse of microglial MHC-I pathway gene expression with aging. To examine the trajectory of MHC-I gene expression, microglial RNA from female Cx3cr1^Litt^ NuTRAP mice from 12-23 months of age was isolated by TRAP and *B2m*, *H2-D1*, *H2-K1*, *H2-M3*, and *H2-Q6* gene expression examined by qPCR. A steady increase in gene expression was observed for all transcripts (One-Way ANOVA with SNK post hoc comparisons (*p<0.05, **p<0.01)).

*Single cell mouse MHC-I protein expression:* To extend these gene expression findings to protein, cell surface MHC-I expression was examined by flow cytometry using cells prepared with our methods optimized to avoid ex vivo activational artifacts in microglia^66^. Antibodies that recognize specific MHC-I β chains are limited but pan-H2 antibodies have been widely used^73^. To validate which brain cells express cell surface MHC-I, hippocampal cells from male and female wild type mice (∼5-7 m.o.) were gated on CD11b^+^ and CD45^mid^ for microglial cells and staining for pan-H2 was assessed (**Figure 4A**). Across all cells less than 10% were pan-H2 positive while ∼95% of CD11b^+^/CD45^mid^ cells, microglia, were pan-H2 positive. Only ∼5% of negative gate cells were pan-H2 positive which indicates that some cells other than prototypical microglia expression MHC-I, potentially neurons, CD45^high^ macrophages, and some microglia as the gating is not 100% efficient. However, this does demonstrate that the principle MHC-I positive cell type in the hippocampus is microglia. To examine whether cell surface MHC-I is altered with aging, hippocampal cells from wild-type 5 m.o. and 24 m.o. male and female mice were gated on CD11b^+^ and CD45^mid^ and again showed enrichment for MHC-I as well as increased fluorescent intensity in old as compared to young, a finding which was observed with two different antibodies (**Figure 4B&C**). Similarly, hippocampal single cell suspensions from Cx3cr1-NuTRAP mice were gated on the GFP^+^ signal and demonstrated increased cell surface expression of MHC-I in 24 m.o. mice as compared to 7-9 m.o. mice (**Figure 4D**).

**Figure 4:**
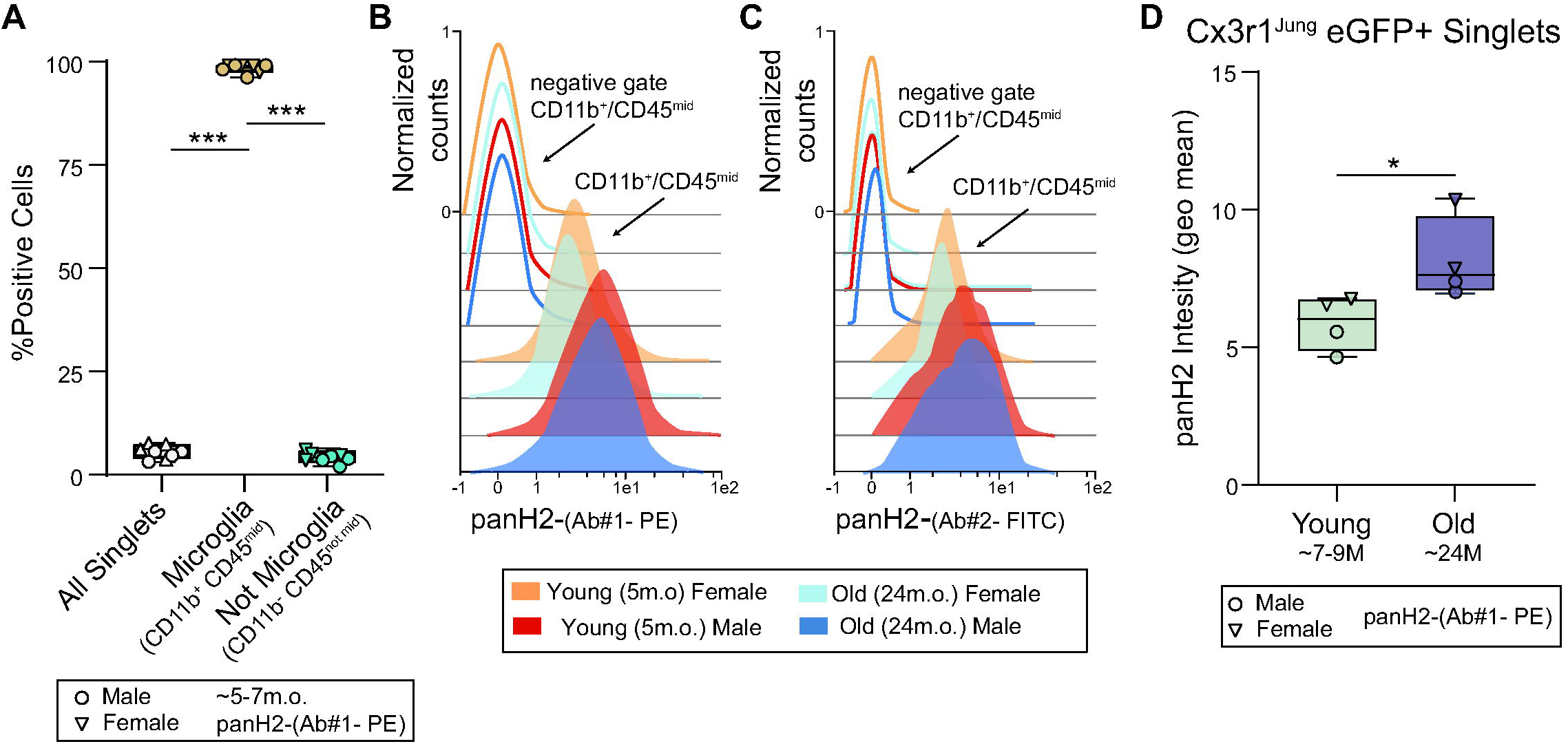
Cell surface microglial MHC-I protein expression increases with aging. **A)** In young (5-7m.o.) mice (n=6) CD11b^+^/CD45^mid^ hippocampal singlets were >95% positive for panH2 staining while the negative gate only ∼5% of cells were positive. One-Way ANOVA with SNK post hoc comparisons (***p<0.001). **B)** In young (5m.o.) and old (24m.o.) males and females panH2 cell surface staining intensity was greater in hippocampal CD11b^+^/CD45^mid^ singlets as compare to the negative gate and was more intense in old as compared young. T-test,*p<0.05, n=4/age. **C)** This finding was recapitulated with a second panH2 antibody. **D)** Male and female, young (5m.o) and old (24m.o.) Cx3cr1^Jung^ NuTRAP mouse hippocampal singlets were gated for EGFP and then assessed for panH2 staining intensity. Staining intensity was greater in old as compared to young mice (t-test, *p<0.05).

To further validate localization of the MHC-I pathway to microglia vs astrocytes we conducted immunohistochemical analysis of brain sections of Aldh1l1-NuTRAP mice where astrocytes can be visualized by specific labeling with the EGFP reporter. Immunolabeling of Aldh1l1-NuTRAP brain tissue sections with CD11b, B2m, and H2Db antibodies showed localization of MHC-I pathway components to microglia and not astrocytes in young (∼3M old) hippocampus (**Figure 5A-B**). Additional immunohistochemical analysis of WT brain sections stained with NeuN, H2Db, and MHCI RT1A antibodies recapitulated the pattern of biased expression of the MHC-I pathway toward microglia, relative to neuronal cells of young (∼3M) hippocampus (**Figure 5C-D**). These qualitative findings were consistent with the quantitative flow cytometry analyses (**Figure 4**) demonstrating that in hippocampus MHC-I pathway is enriched, almost exclusively, in microglia. Lastly, to further validate microglial induction with aging, B2m co-localization in hippocampal sections from young (6 m.o.) and old (21 m.o.) Cx3cr1-NuTRAP mice demonstrated B2m immunoreactivity in EGFP^+^ CD11b^+^ cells and qualitative increases with aging (**Figure 5E**).

**Figure 5:**
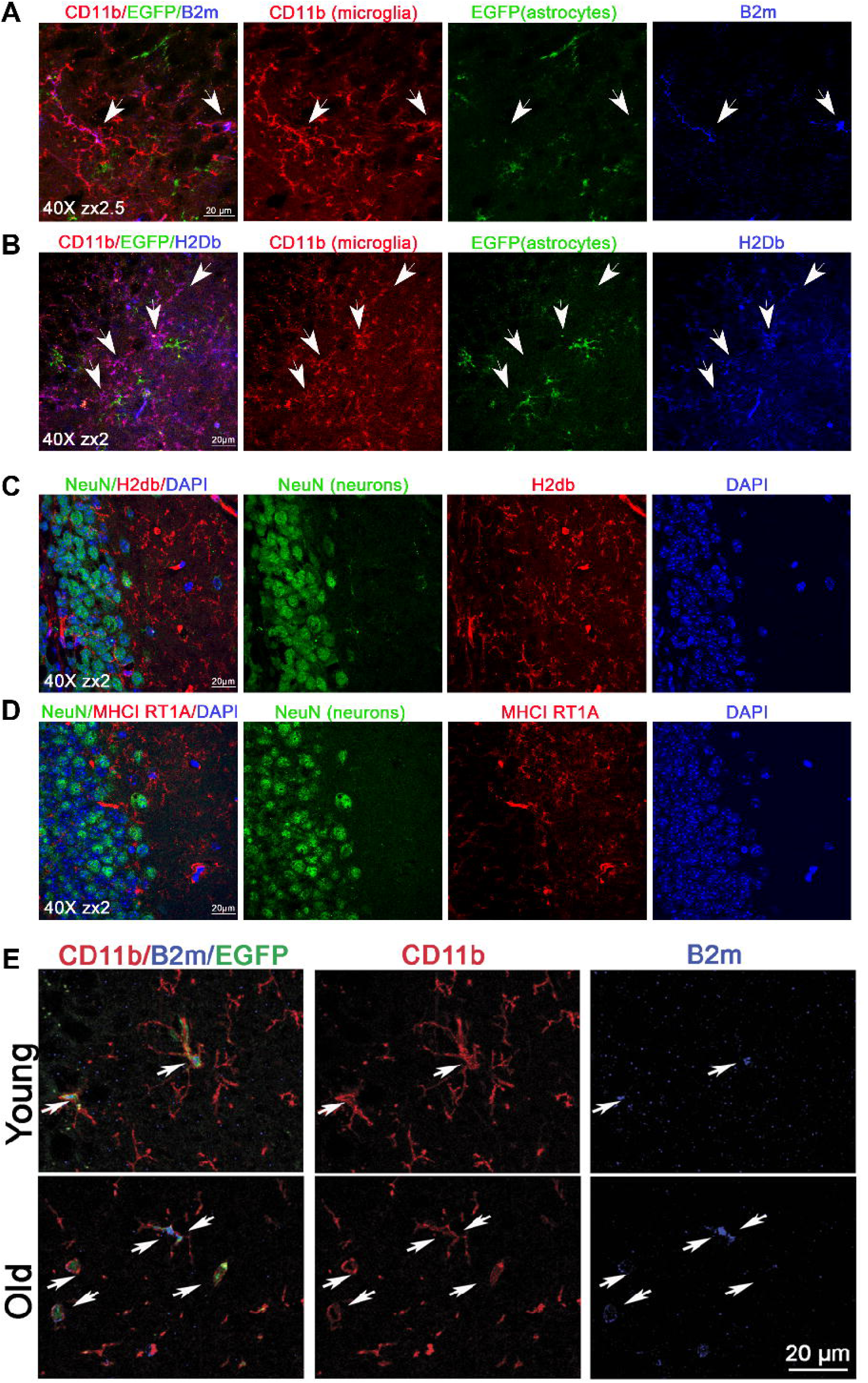
Microglial MHC-I cellular localization and B2m induction with aging. In hippocampal sections from Aldh1l1-NuTRAP mice, astrocytes were visualized the EGFP reporter. Immunolabeling with CD11b and either B2m (**A**) or H2Db (**B**) antibodies showed localization of MHC-I pathway components to microglia and not astrocytes in young (∼3M old) hippocampus. WT brain sections were stained with NeuN for neurons and H2Db (**C**) or MHCI RT1A (**D**) antibodies showed no co-localization. In hippocampal sections young, (6 m.o.) and old (21 m.o.) Cx3cr1^Jung^-NuTRAP mice stained for CD11b and B2m demonstrated qualitative induction of B2m staining in CD11b^+^/GFP^+^ cells (**E**).

*MHC-I expression in human CNS cell type:* To assess cell type-specific MHC-I expression and regulation with aging in humans expression data from public repositories was collected and re-analyzed. The Genotype-Tissue Expression Project (GTeX)^74^ dataset was collected with age and tissue annotations. MHC-I components were assessed as a function of age in blood, brain, liver, lung, and muscle samples. A consistent induction with aging was observed in brain, lung, and muscle but not in blood or liver annotated samples (**Figure 6A**). RNA-Seq data from sorted human brain cells^75^ was assessed for MHC-I expression and while *B2m* was expressed across all examined cell types, other MHC-I components were almost exclusively expressed in microglia (**Figure 6B, Supplemental Table 7**). Microglial RNA-Seq data collected from adult (average age 53 y.o.) and aged (average age 93 y.o.) fresh brain samples isolated by magnetic bead enrichment for CD11b+ cells followed by FACS for CD11b^+^/CD45^+^^76^ was next assessed. MHC-I components *B2m* and canonical and non-canonical MHC-I, were induced with aging (**Figure 6C**).

**Figure 6:**
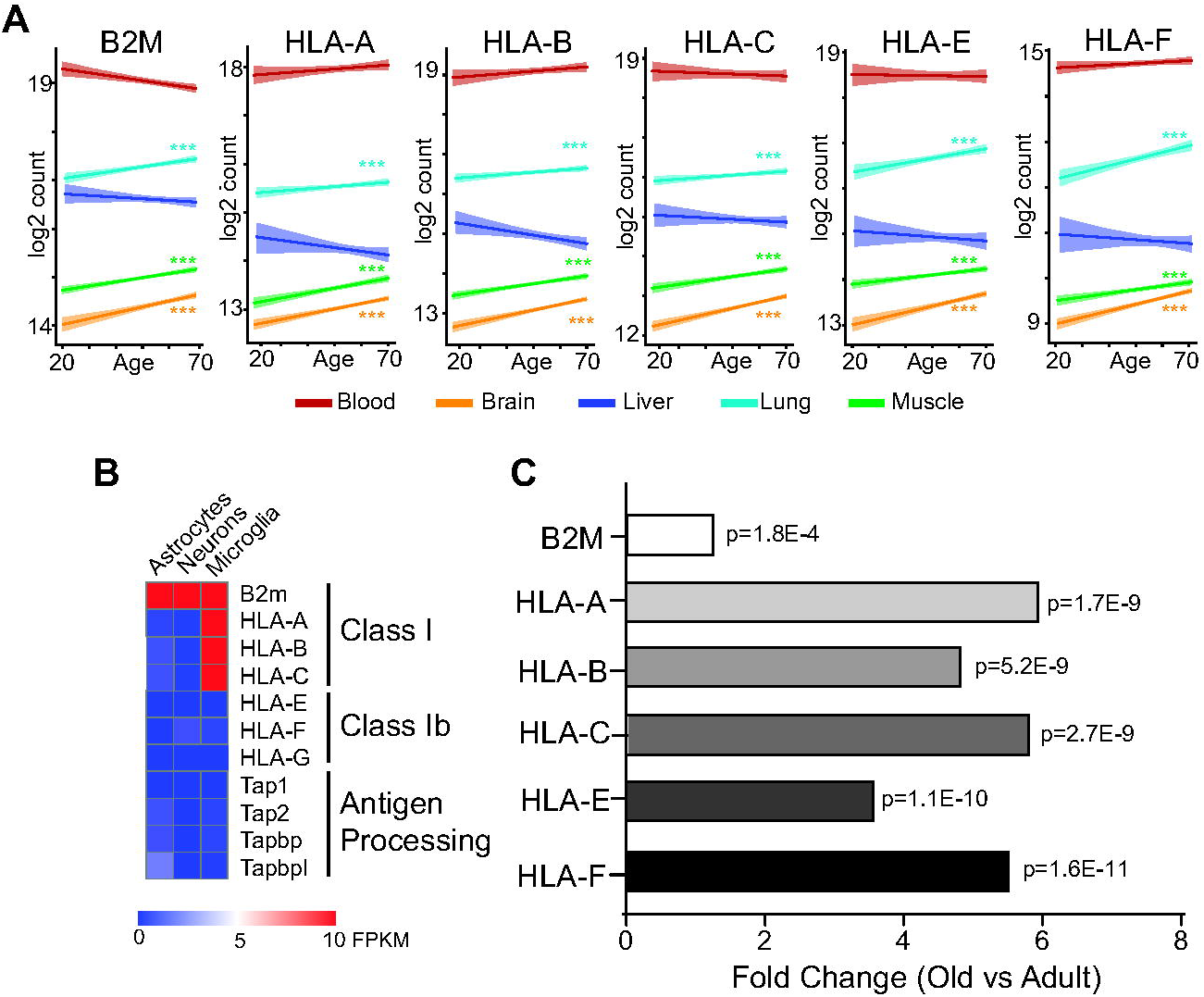
MHC-I pathway expression is induced in human microglia with brain aging. **A)** In re-analysis of whole tissue data from the human GTEx database, induction of alpha chain MHCIs *HLA-A*, *-B*, *-C*, *–E*, and *-F* and the invariant beta chain (*B2m*) is observed in brain, and muscle and lung, but not in circulating blood cells or liver. Pearson’s r, ***p<0.001. **B)** Examination of human transcriptome patterns by cell nuclei sorting reveals the MHC-I pathway is predominantly expressed by microglia. **C)** In human microglia expression of the MHCI pathway is elevated with aging. n=5 per group, Adult = Ave. 53y.o. Aged = Ave. 93 y.o. T-test.

*Single cell co-expression analysis in mouse brain*: To examine co-expression of MHC-I pathway components at the individual cell level in the mouse brain, the Tabula Muris^67^ and the Atlas of the Aging Mouse Brain^68^ were retrieved and analyzed. Almost all individual microglia expressed both the invariant β chain (*B2m*) and at least one α chain needed to make a functional, cell surface MHC-I (**Figure 7A**). Very limited numbers of astrocytes or neurons were found to express both α and β chains required for a functional MHC (**Figure 7B&C**). During these analyses it was observed that many microglial also expressed paired immunoglobulin-like receptors (*Lilra5*, *Lilrb4*, *Pilra*, *Lilrb3* (aka, *PirB*)), which are potential receptors for MHC-I^42^. No expression of these receptors was observed in astrocytes or neurons. This could allow for microglial MHC-I signaling in a cell autonomous manner and between microglia in cell non-autonomous manner. Therefore, the expression of paired immunoglobulin-like receptors across cell types and with aging was examined.

**Figure 7:**
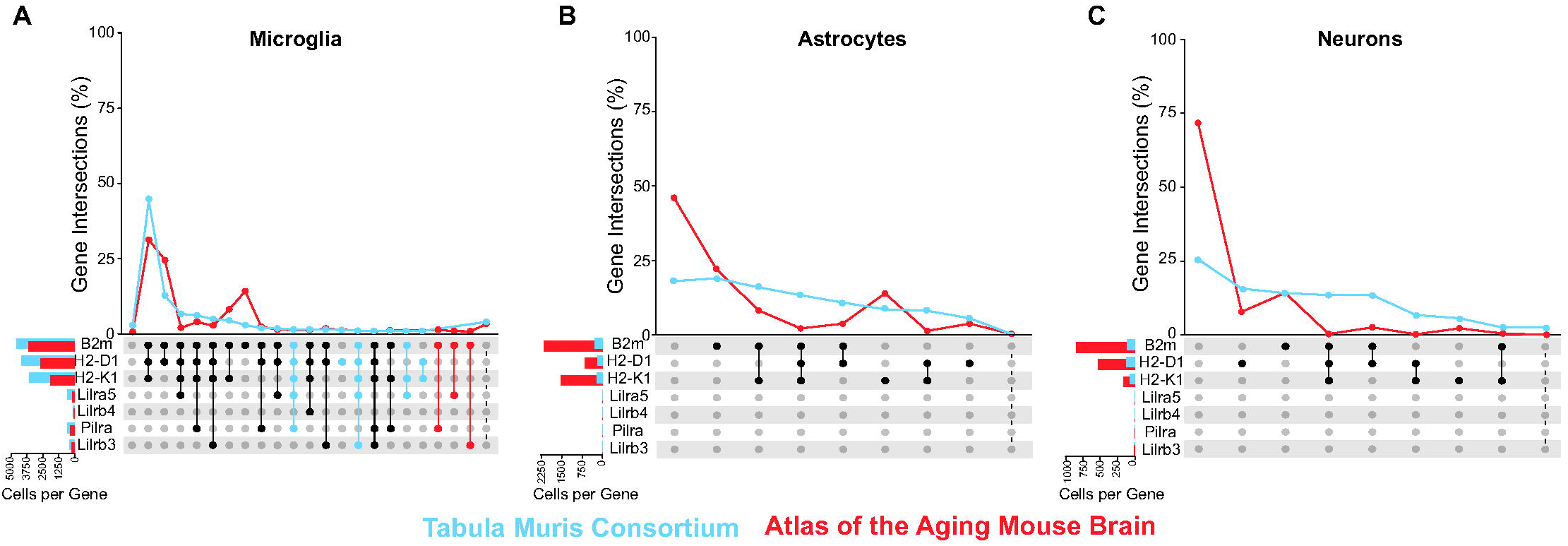
Single cell expression of MHC-I pathway components and receptors. Using public atlas data from the Tabula Muris and Atlas of the Aging Mouse Brain, individual cell co-expression of MHC-I pathway components and potential receptors was examined in **A)** microglia, **B)** astrocytes, and **C)** neurons. In microglia, nearly all cells express MHC-I α and β chains while in astrocytes and neurons only a small portion of cells expressed both components. Additionally potential co-expression of MHC-I receptors *Lilra5*, *Lilrb4*, *Pilra*, and *Lilrb3* was examined. A significant portion of microglia demonstrated co-expression of at least one receptor while no receptor expression was observed in astrocytes and neurons. Black circles in the upset plot represent patterns observed in both studies while blue or red circles are patterns only observed in one study.

*Cellular localization and expression of mouse Pilr and Lilr genes with aging:* With the finding that Pilr and Lilr genes are co-expressed with MHC-I in microglia, the cell type-specific expression of these genes and their regulation with age was asssed. In the same cell sorting and translatome datasets as **Figure 1**, Pilr and Lilr receptor family expression was restricted to microglia (**Figure 8A, Supplemental Table 3**). Across microglial, astrocyte, and neuronal TRAP-qPCR aging data, receptor expression of *Lilrb3*, *Lilrb4*, and *Pilra* were elevated with aging in microglia but not astrocytes or neurons (**Figure 8B-D, Supplemental Table 5**). In fact, most of these receptors were not detected in TRAP isolated RNA from astrocytes and neurons. Extending to a timecourse of microglial aging, *Lilrb3* and *Lilrb4* increased at 18-21 m.o. as measured by TRAP-qPCR, similarly to the MHC-I α and β chains (**Figure 8E, Supplemental Table 6**).

**Figure 8:**
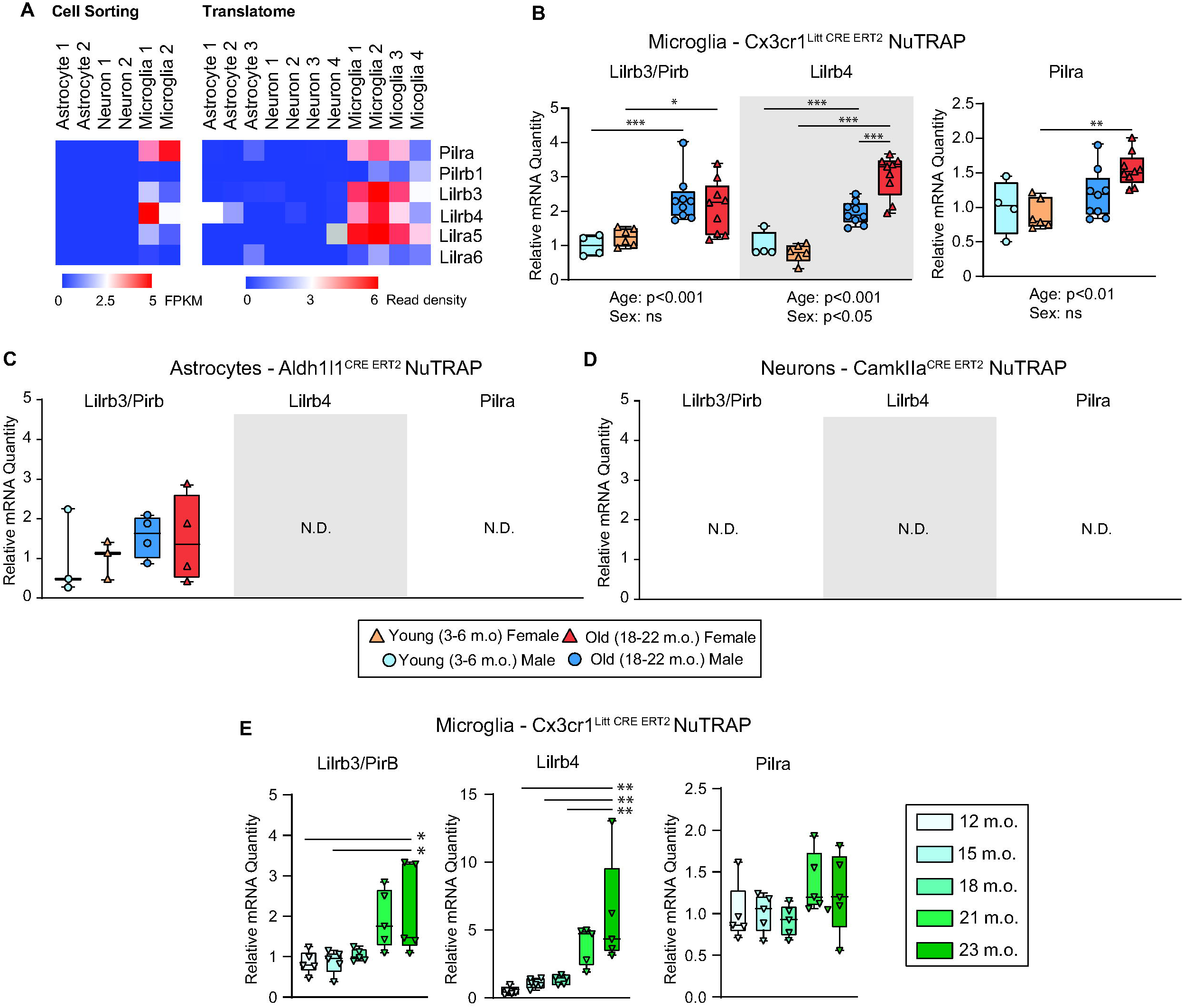
Cell type specific MHC-I receptor expression. **A)** Using bulk RNA-Seq data from sorted CNS cell types and from TRAP-Seq data of cell type-specific NuTRAP mice lines MHC-I receptors were nearly solely expressed by microglia. **B-D)** Cell type-specific hippocampal RNA from microglial (Cx3cr1^Litt^), astrocyte (Aldh1l1), and neuronal (CamkIIa) NuTRAP lines was isolated by TRAP. qPCR analysis was performed on Young (3-6M) and Old (18-22M) male and female mice. Increased microglial levels of *Lilrb3* (also known as *PirB*), *Lilrb4*, and *Pilra* were observed in microglia while no changes or no expression were observed in astrocytes and neurons (Two-Way ANOVA (Factors: Age and Sex) with SNK post hoc comparisons (*p<0.05, **p<0.01, ***p<0.001)). **E)** To examine the trajectory of MHC-I gene expression, microglial RNA from female Cx3cr1^Litt^ NuTRAP mice from 12-23 months of age was isolated by TRAP and *Lilrb3*, *Lilrb4*, and *Pilra* gene expression examined by qPCR. A steady increase in *Lilrb3* and *Lilrb4* gene expression was observed for all transcripts (One-Way ANOVA with SNK post hoc comparisons (*p<0.05, **p<0.01)).

*Cellular localization of human Pilr and Lilr genes and expression with aging:* In human GTeX data, Lilr and Pilr family gene expression increased as a function of age in brain with a variety of patterns observed in other tissues (**Figure 9A**). Cell sorting data from human brains showed, similarly to mice, restricted expression of these receptors to microglia (**Figure 9B**), with the exception of *Pilrb*. In human microglia, Lilr and Pilr family gene expression increased with age (**Figure 9C**). Thus, microglial enriched expression of Pilr and Lilr gene family members is conserved in mice and humans and increases with age.

**Figure 9:**
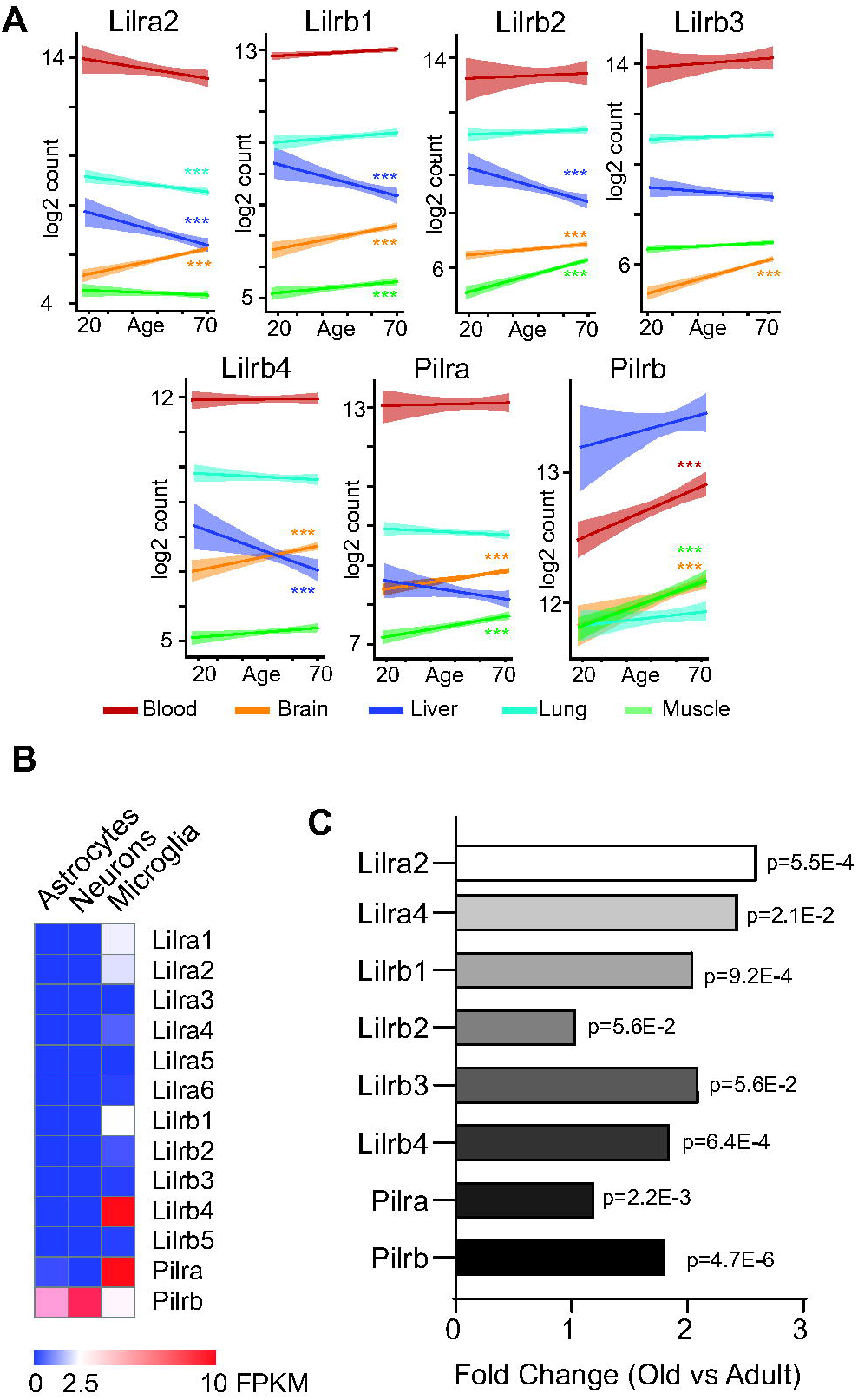
MHC-I receptors are induced in human microglia with brain aging. **A)** In re-analysis of whole tissue data from the human GTEx database, induction of alpha chain MHC-I receptors were increased in the brain with aging and a variety of patterns in other tissues. Pearson’s r, ***p<0.001. **B)** Examination of human transcriptome patterns by cell nuclei sorting reveals MHC-I receptors are predominantly expressed by microglia. **C)** In human microglia expression of the MHC-I receptors is elevated with aging. n=5 per group, Adult = Ave. Age 53 Aged = Ave. Age 93. T-test.

*Microglial MHC-I pathway regulation in mouse models of AD and human samples*: AD is one of the most common and debilitating age-related neurodegenerations. To extend these findings of age-related microglial MHC-I and paired immunoglobulin-like receptor induction, a variety of AD mouse models (rTg4510^48^, AppNL-G-F^48, 77^, App-SAA (GSE158153) and 5xFAD^78, 79^) with microglial-specific expression data were examined for altered expression of MHC-I and paired immunoglobulin-like receptors. A consistent microglial up-regulation of almost all the genes examined was evident, across models and technical methods with differing degrees of cell purity (single-cell>FACS>antibody bound magnetic beads) and different levels of sensitivity for detecting gene expression (bulk>sc/snRNA-Seq) (**Table 1**). The reproducibility of upregulation across models and methods increases confidence in the findings.

**Table 1:**
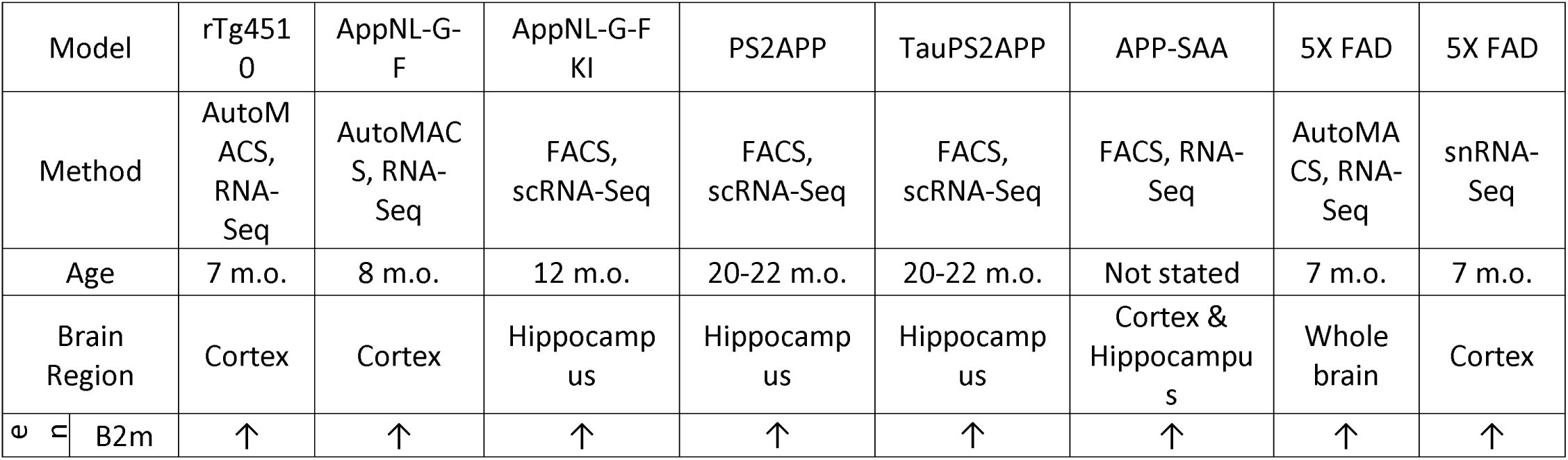
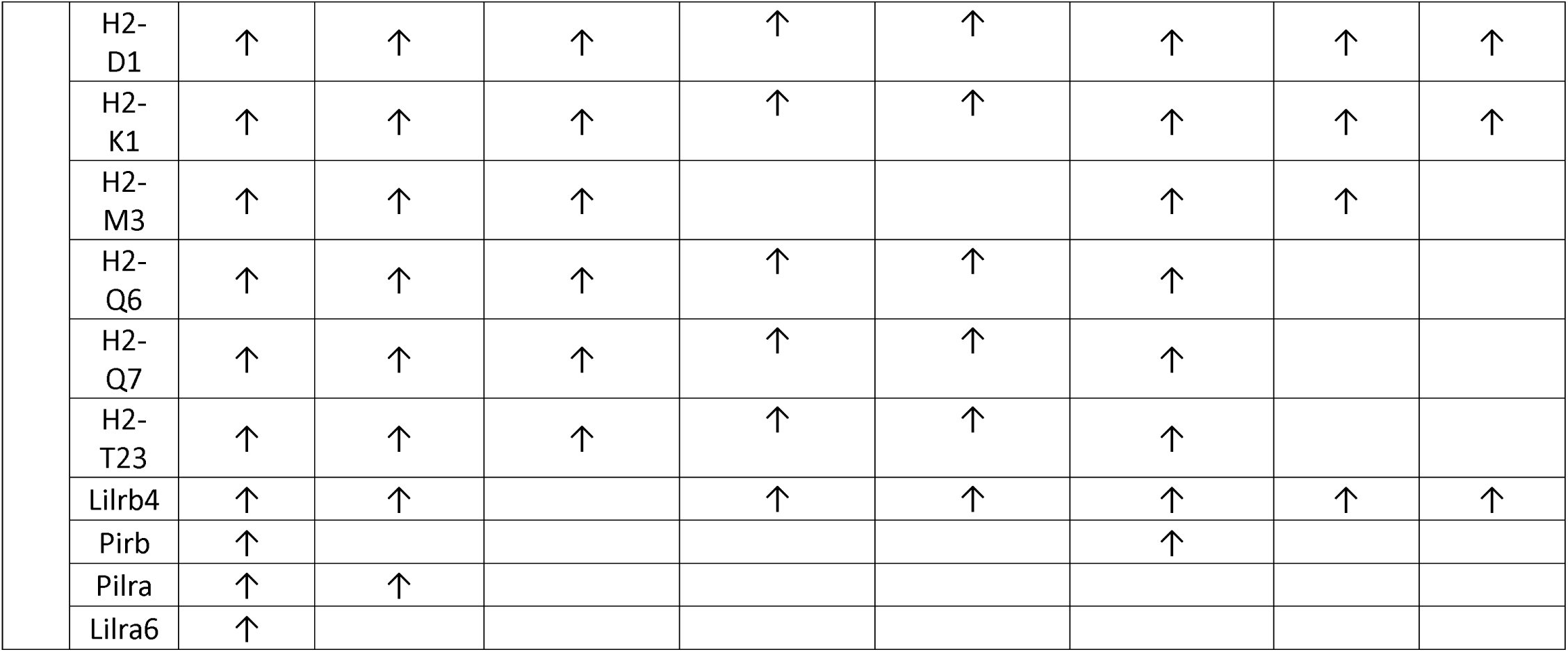
Alterations in MHC-I pathway components and paired immunoglobulin-like receptors in mouse models of AD.

Building on these results we isolated microglia from 12 m.o. APP-PSEN1 hippocampus by CD11b^+^ microbeads and performed RNA-Seq. The transcriptomic patterns of APP-PSEN1 microglia differed from controls (**Figure 10A**) and 4,440 genes demonstrated differential expression (t-test, Benjamini-Hochberg Multiple Testing Correction, FC>|1.25|). (**Figure 10B, Supplemental Table 8**). To assess whether the microglial expression patterns may be indicative of shifts in phenotypic subtypes, gene sets were developed for Activated Response Microglia (ARM)^77^, Axon Tract Microglia (ATM)^80^, Disease Associated Microglia (DAM)^81^, Interferon Response Microgla (IRM)^77^, Lipid Droplet Accumulating Microglia (LDAM)^82^, Neurodegenerative Microglia (MGnD)^49^, Proliferative Region Associated Microglia (PAM)^83^, and White Matter Associated Microglia (WAM)^84^ as well as homeostatic^66^ and macrophage marker lists^85–87^(**Supplemental Table 9**). Both z-scores (**Figure 10C**) and GSEA analysis (**Figure 10D**) were performed with these lists and a shift in favor of PAM, DAM and WAM phenotypes were evident in both analyses. As well, homeostatic markers were depleted and notably LDAM markers were suppressed. For the MHC-I genes of interest, an upregulation of *B2m*, *H2-D1*, *H2-K1*, and *H2-M3* was observed, as was an induction of *Lilrb4* and intriguingly, a suppression of *Pilra* (**Figure 10E**). In parallel we examined human AD microglial data^79, 88, 89^ from tissue, sorted cells or snRNA-Seq data and again found induction of MHC-I and paired immunoglobulin-like receptors in cases as compared to controls (**Table 2**).

**Figure 10:**
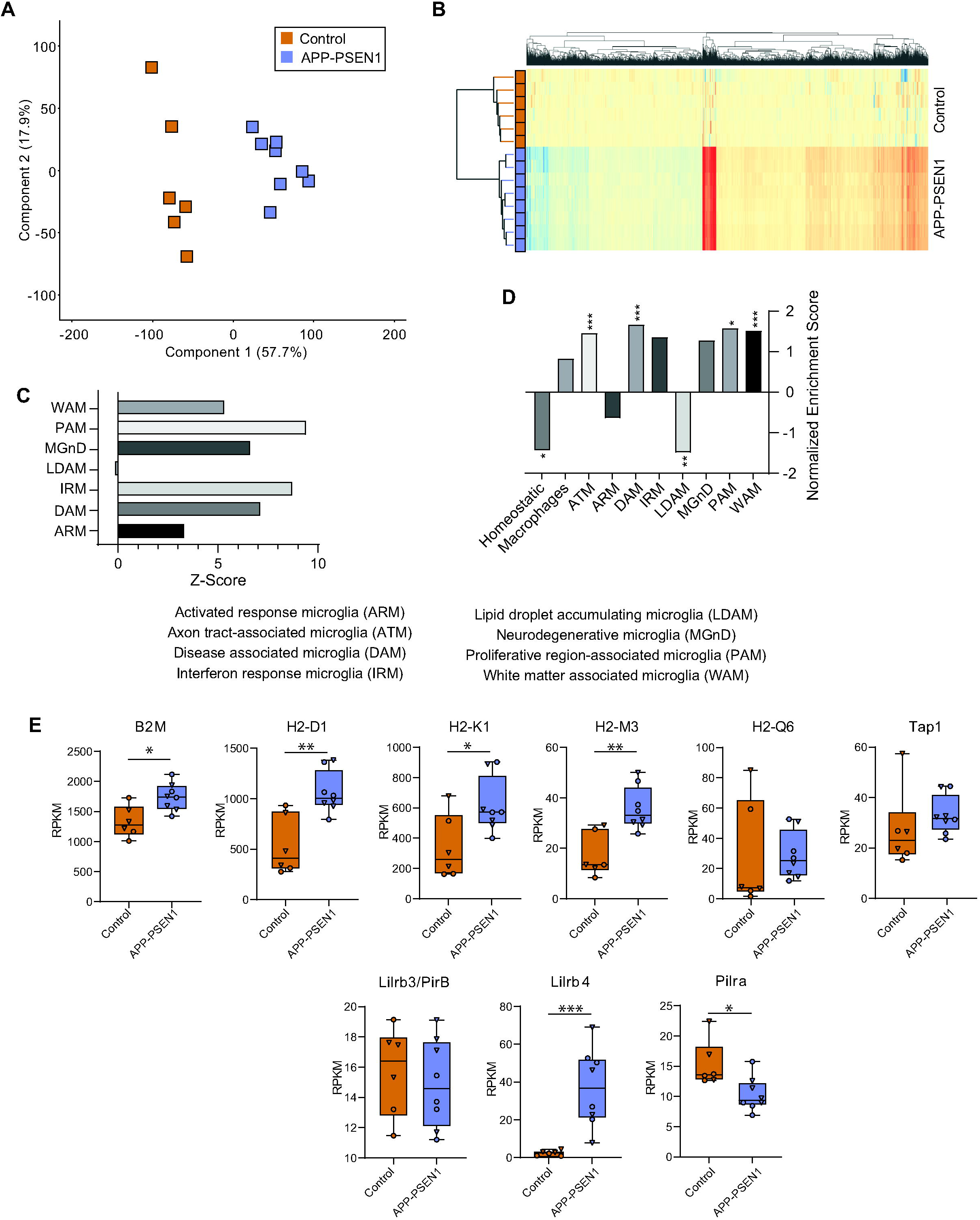
APP-PSEN1 hippocampal microglial transcriptome. **A)** Whole transcriptome pattern of gene expression in CD11b+ hippocampal cells separates between 12 m.o. control and APP-PSEN1 mice. **B)** Clustering of 4,389 differentially expressed genes (t-test, B-H MTC, FC>|1.25|). **C)** Using previously described microglial phenotypic gene expression patterns activation and repression z-scores were generated, indicating expression patterns consistent with increased presence of WAMs, PAMs, MgnDs, IRMs, DAMs, and ARMs. **D)** Gene Set Enrichment Analysis (GSEA) of the non-overlapping phenotypic gene expression sets revealed a similar increased enrichment of ATMs, DAMs, PAMs, and WAMs, while homeostatic patterns were suppressed in APP-PSEN1 as compared to control mice (nominal p-value, *p<0.05, **p<0.01, ***p<0.001). **E)** Examination of specific MHC-I pathway components reveals increased expression of most components in APP-PSEN1 microglia. T-test, *p<0.05, **p<0.01, *** p<0.001.

**Table 2:** Alterations in MHC-I pathway components and paired immunoglobulin-like receptors in human AD.

| Study |  | The Mount Sinai Brain Bank (MSBB) study | Cribbs et al., | Rush Alzheimer's Disease Center |
| --- | --- | --- | --- | --- |
| Method |  | RNA-Seq | Microarray | snRNA-Seq |
| Brain Region |  | Parahippocampal gyrus | Hippocampus | Prefrontal cortex |
| Gene | *B2m* | ↑ |  | ↑ |
|  | *HLA-A* | ↑ |  | ↑ |
|  | *HLA-B* | ↑ | ↑ | ↑ |
|  | *HLA-C* | ↑ | ↑ |  |
|  | *HLA-E* | ↑ | ↑ | ↑ |
|  | *HLA-F* |  |  |  |
|  | *HLA-G* |  | ↑ |  |
|  | *Lilrb3/PirB* | ↑ |  | ↑ |
|  | *Lilrb4* | ↑ |  | ↑ |
|  | *Pilra* | ↑ |  |  |

*Senescent marker p16INK4A microglial expression parallels MHC-I induction*: Recent findings in fibroblasts with aging demonstrate that MHC-I induction may be a mechanism by which senescent cells avoid clearance^90^. An observation from the APP-PSEN1 RNA-Seq data was an increase in the expression level of the senescence marker *p16INK4A*, but not the other *Cdkn2a* transcript *p19ARF* in microglia (**Figure 11A**). These two isoforms can be discriminated by the unique 5’ exons of the two transcripts (**Figure 11B**) (something not observable in most single-cell RNA sequencing studies). TRAP-qPCR was performed using the aging samples from each cell type described above using a *p16INK4A*-specific qPCR assay. A significant induction of *p16INK4A* with aging was observed in microglia but no expression was detected in astrocytes or neurons (**Figure 11C**). In microglial timecourse samples, p16INK4A expression followed a similar pattern as the MHC-I genes, with a gradual increase that accelerated at 21 m.o. (**Figure 11D**). The similar patterns of expression across the time course study was evident in a correlation analysis with high degrees of correlation across genes examined (**Figure 11E**).

**Figure 11:**
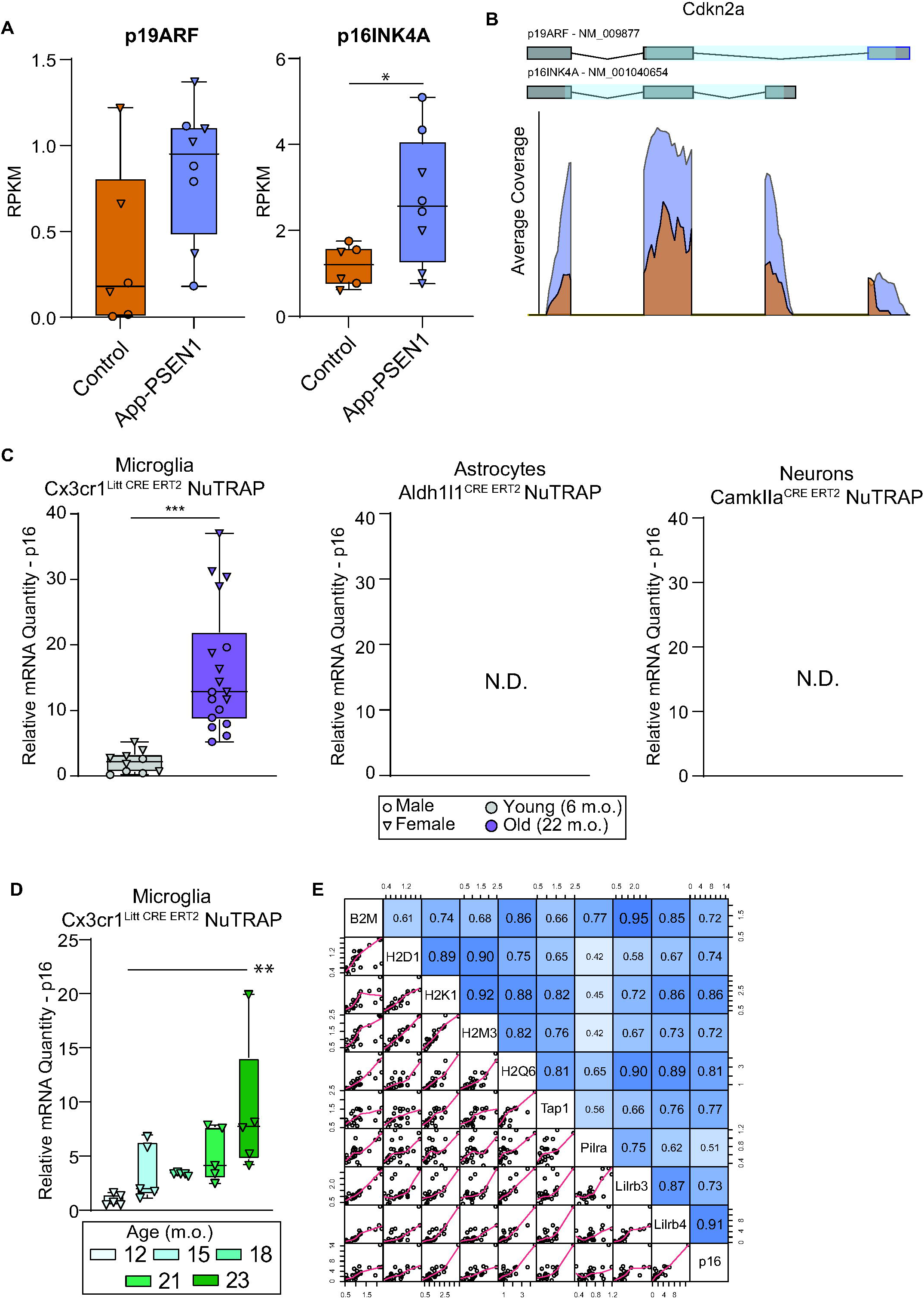
Aging and Alzheimer’s models are associated with increased microglial expression of senescence marker p16INK4a. **A)** In CD11b+ hippocampal cells from 12 m.o. control and APP-PSEN1 mice transcript-specific analysis of *Cdkn2a* revealed increased expression of *p16INK4A* but not *p19ARF*. **B**) Differentiation of *p16INK4A* from *p19ARF* is possible by alternate 5’ exons in the two transcripts. **C**) Cell type-specific hippocampal RNA from microglial (Cx3cr1^Litt^), astrocyte (Aldh1l1), and neuronal (CamkIIa) NuTRAP lines was isolated by TRAP. qPCR analysis was performed on Young (3-6M) and Old (18-22M) male and female mice for *p16INK4A*. Increased microglial levels of *p16INK4A* were observed and no detectable *p16INK4A* transcript was observed in astrocytes and neurons (t-test, ***p<0.001). **D**) Microglial *p16INK4A* expression as measured by qPCR increased steadily from 12M to 23M of age. One-Way ANOVA with SNK post hoc comparisons (**p<0.01). E) MHC-I pathway components, putative receptors, and *p16INK4A* all positively correlated across the 12-23M data.

*MHC-I pathways genes a potential AD intervention targets*: The common regulation of these MHC-I pathway genes in microglia across aging and AD, and in mouse and man, provides a compelling result that this is a conserved phenomenon. To further examine if these genes warrant further mechanistic study, TREAT-AD scores^91^ were collected for each of these genes and compared to the distribution of all annotated genes (**Figure 12**). The TREAT-AD score was developed as part of NIH consortia to prioritize AD targets for investigation and potentially therapeutic development. The TREAT-AD score is a combination of genetic associations, predicted variant impacts, and associated dementia phenotypes, paired with extensive sets of transcriptomic and proteomic AD-related expression changes. A notable rightward skew of MHC-I pathways and paired receptors suggests this pathway may warrant further investigation.

**Figure 12:**
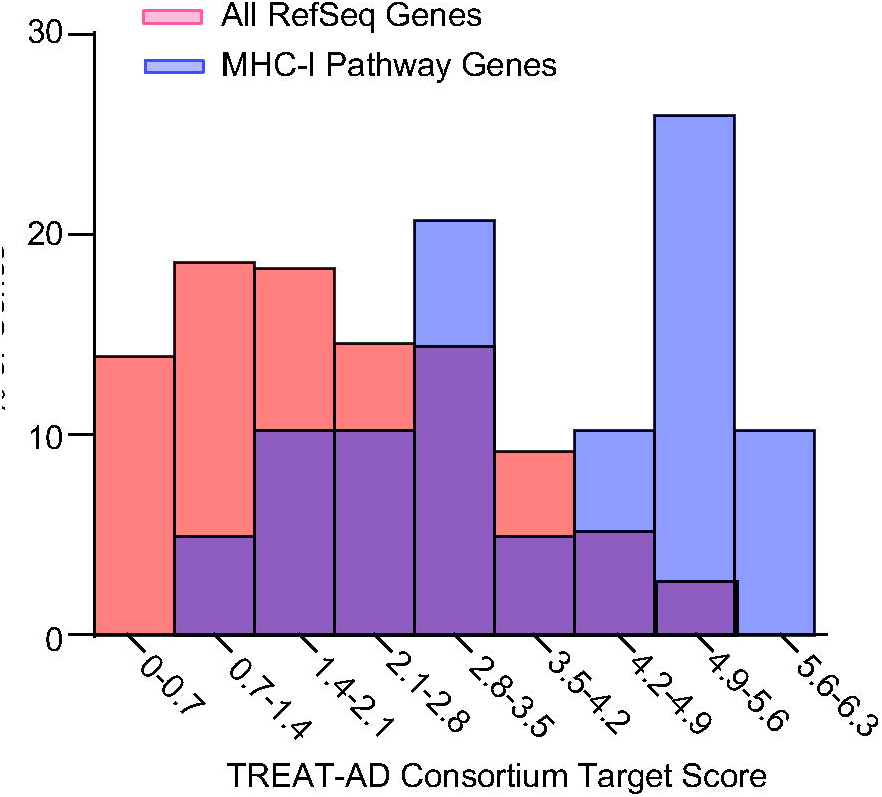
MHC-I pathway is enriched for high TREAT-AD target scores. TREAT-AD scores are derived from genetic associations, predicted variant impact, and linkage with dementia-associated phenotypes as well as transcriptomic and proteomic data studies to rank genes for potential investigation in AD studies. MHC-I α and β chain genes and potential non-TCR receptors were compared to all RefSeq genes and found to be over-represented for high scores, which are indicative of higher priority AD targets. By way of comparison well known targets have scores of 6.05 – *Trem2*, 5.56 – *Apoe*, and 5.05 – *Tyrobp*.

## Discussion

These results demonstrate that microglia are the principal site of MHC-I expression in the CNS, and that with aging and in human AD samples or in mouse models of AD, microglial MHC-I increases in expression at the transcript, translating transcript, and protein level. This provides a cellular context to previous tissue level analyses with aging^52, 53^. The consistency of these finds across species, studies, technical methods, and microglial isolation approaches gives robustness to the findings and could serve as the basis for future studies to understand the function of microglial MHC-I in health and disease. Other cell types in the CNS, i.e., neurons, may or are even likely to also express MHC-I^43, 92, 93^ but from the data presented here this expression is at a much lower level in health and disease. The finding of microglial as the principle source of MHC-I in the CNS also gives context to previous constitutive mouse knockout studies that were interpreted with solely a neuronal focus and may need to be reinterpreted^43^.

The functional consequences, if any, of microglial MHC-I induction with aging and AD remain to be determined. The canonical signaling partner for MHC-I is the TCR. The potential for T-cell infiltration into the brain with aging and AD has been a question in the field for at least a decade^94, 95^. Aging causes an increase in CD8+ T-cells in mice^96^, which has subsequently been characterized as tissue resident T-cells (Trm), but the characterization of infiltrating T-cells with aging and AD is an ongoing and controversial topic (for review see^95, 97–99^). The data on T-cell infiltration into the parenchyma with non-pathological aging is varied, with reports ranging from no infiltration to significant numbers^100–102^. While this question remains to be definitively answered, microglial reactivation could serve to recruit T-cells^103^, and microglial proximity to pericytes and endothelial cells suggest they could present MHC-I to infiltrating T-cells^104^. In turn, T-cells could modulate microglial phenotype^105^. In the APP-PSEN1 model used here, infiltration of T-cells has been reported^106^, which means one possibility is that the induction of MHC-I observed here could be to signal to T-cells. Antigens can exit the brain through the dural sinus^107^ for positive selection, but whether this occurs with aging or AD^108^ is still being determined. Nonetheless, the possibility exists for antigen-specific signaling of microglial MHC-I to T-Cells with aging and AD, though much further investigation is needed.

The second principal finding from this study is that antigen-independent MHC-I receptors – leukocyte immunoglobulin-like receptor subfamily receptors (Lilrs)^31, 32^ and paired immunoglobulin-like type 2 receptors (Pilrs)^33^ – are present and induced in microglia with aging and AD. These receptors are almost exclusively restricted to microglia in these data. These receptors contain ITAMs (Immunoreceptor Tyrosine-based Activation Motifs) and ITIMs (Immunoreceptor Tyrosine-based Inhibition Motifs) that can induce activational or inhibitory signals in the recognizing cell^35^. ITAMs are familiar to AD research through Trem2 effector Tyrobp/DAP12 which contains an ITAM motif^109^. ITAMs activate and ITIMs inhibit Syk activity, a central controller of microglial phenotype^110–112^. Lilr and Pilr family receptors can bind MHC-I^32, 113^ in antigen-dependent and antigen-independent manners and could engage ITAM or ITIM cascades and modulate Syk activity depending on the receptor. MHC-I signaling through these receptors requires a great deal more basic investigation to fully elucidate this process^28–30^. With both MHC-I and Lilr and Pilr family receptors being present on microglia, MHC-I could be signaling in a cell-autonomous manner (or cell non-autonomously in microglia-microglia contacts) to regulate microglial phenotype through these receptors, ITIMs/ITAMs and Syk. This concept is supported by data such as the ITIM containing CD22 receptor regulating microglial phagocytosis^114^. This area of investigation clearly warrants further investigation given the consistent findings in humans and mouse models with aging and neurodegeneration.

Our microglial hypothesis differs but is not in opposition to the neuron-centric MHC-I function in AD recently reported^43^ in that MHC-I, driven by ApoE expression, drives neurodegeneration. As shown in that report, ApoE expression is highest in microglia and astrocytes with a small minority of neurons expressing low levels of ApoE. Global knock-in of human ApoE4 and then knockout of ApoE specifically in neurons reduced expression of some MHC-I genes (*H2-T22*, *H2T23*, *H2-T24*, and *H2-D1*, but not *H2-K1*). We propose that their studies showing protection from p-tau in a non-cell-specific *B2m* knockout model are actually mediated through microglia instead of neurons, as the vast majority of B2m is in microglia. Taking our cell type specific TRAP-Seq data across ages and sexes, we find that *ApoE* levels are highly correlated to MHC-I components *B2m*, *H2-D1*, and *H2-K1* in microglia and neurons, but the expression level is an order of magnitude higher in microglia than neurons (data not shown). Thus, we would propose a microglial focus to MHC-I actions in brain aging and AD where we have found robust and consistent upregulation of MHC-I and potential receptors.

Lastly the finding of the canonical senescence marker, p16INK4A, as also induced in microglia, but not other cell types examined, could point to another function of microglial MHC-I. In the periphery, MHC-I expression in senescent fibroblasts has been demonstrated to be a mechanism by which senescent cells avoid immune clearance through the ITIM containing receptor NKG2A on NK and T cells^90^. Microglial senescence has been described^115^ and is attributed to replicative exhaustion^116^. In fact, an analysis of p16INK4A^+^ versus p16INK4A-microglia show higher levels of the MHC-I, Lilr, and Pilr genes observed here^117^. The nature of microglial ‘senescence’^118^ still requires more study, and it remains to be determined if MHC-I induction is a result or cause of microglial senescence, or if senescence is even the best categorization of this phenotypic state of microglia.

In conclusion these data show that microglia express MHC-I at much higher levels than other CNS cell types examined in mice and man. Furthermore, MHC-I is induced in microglia with aging and AD (or AD models) in humans and mice and occurs concurrently with induction of the senescence marker p16INK4A. While the signaling partner(s) of MHC-I are unknown, we report that microglia co-express Lilr and Pilr family receptors that could bind MHC-I to transduce inhibitory or excitatory signals to microglia. Intriguingly, these receptors are also induced in microglia with aging and AD, identifying a potential new pathway of microglial phenotype regulation that may also be associated with microglial senescence. Microglial specific manipulations of MHC-I in animal models with aging and in AD models are needed to mechanistically understand the role of MHC-I, if any, in the regulation of microglial function in health and disease.

## Declarations

**Acknowledgments**: The Mount Sinai Brain Bank (MSBB) study data published here were data obtained from the AD Knowledge Portal (https://adknowledgeportal.org/). These data were generated from postmortem brain tissue collected through the Mount Sinai VA Medical Center Brain Bank and were provided by Dr. Eric Schadt from Mount Sinai School of Medicine. The authors would like to acknowledge Jay Hicks for assistance with figure preparation.

**Sources of Funding**: The content is solely the responsibility of the authors and does not necessarily represent the official views of the National Institutes of Health. This work was also supported by grants from the National Institutes of Health (NIH) P30AG050911, R01AG059430, DP5OD033443, T32AG052363, F31AG064861, R21AG072811, and F31AG079620. This work was also supported in part by awards I01BX003906, I01BX005396, IK6BX006033, and ISIBX004797 from the United States (U.S.) Department of Veterans Affairs, Biomedical Laboratory Research and Development Service. The content is solely the responsibility of the authors and does not necessarily represent the official views of the National Institutes of Health. The views expressed in this article are those of the authors and do not necessarily reflect the position or policy of the Department of Veterans Affairs or the United States government.

Conflicts of interests/Competing interests: The authors declare they have no financial or non-financial interests directly or indirectly related to this work.

## Notes

### Competing Interest Statement

The authors have declared no competing interest.

### Summary of Updates

Updated text and figures

